# Plasticenta: Microplastics in Human Placenta

**DOI:** 10.1101/2020.07.15.198325

**Authors:** Antonio Ragusa, Alessandro Svelato, Criselda Santacroce, Piera Catalano, Valentina Notarstefano, Oliana Carnevali, Fabrizio Papa, Mauro Ciro Antonio Rongioletti, Federico Baiocco, Simonetta Draghi, Elisabetta D’Amore, Denise Rinaldo, Maria Matta, Elisabetta Giorgini

## Abstract

Microplastics are particles smaller than five millimetres obtained from the degradation of plastic objects abandoned in the environment. Microplastics can move from the environment to living organisms and, in fact, they have been found in fishes and mammals.

Six human placentas, prospectively collected from consenting women with uneventful pregnancies, were analyzed by Raman Microspectroscopy to evaluate the presence of microparticles. Detected microparticles were characterized in terms of morphology and chemical composition.

12 microparticles, ranging from 5 to 10 μm in size, were found in 4 out of 6 placentas: 5 in the foetal side, 4 in the maternal side and 3 in the chorioamniotic membranes. All the analyzed microparticles were pigmented: three of them were identified as stained polypropylene, while for the other nine it was possible to identify only the pigments, which are all used for man-made coatings, paints and dyes.

Here we show, for the first time, the presence of microparticles and microplastics in human placenta. This sheds new light on the impact of plastic on human health. Microparticles and microplastics in the placenta, together with the endocrine disruptors transported by them, could have long-term effects on human health.

## INTRODUCTION

> *“…avoiding the use of plastic and paper, reducing water consumption, separating refuse…”*
>
> — HOLY FATHER FRANCIS From: ENCYCLICAL LETTER LAUDATO SI’ ON CARE FOR OUR COMMON HOME

In the last century, the global production of plastics has grown exponentially by over 350 millions of tons per year, and a part ends up polluting the environment^1^. It has been estimated that 8.3 billion tons of plastic have been produced since the 1950s, with a constant increase in the last three decades. Global production of plastics currently exceeds 320 million tons (Mt) per year, and over 40% is used as single-use packaging, hence producing plastic waste. In Europe, 26 million tons of plastic waste are produced every year; only 30% is collected for recycling, while the rest is burned or ends up in landfills and it is dispersed into the environment. The degradation that plastics undergo when released into the environment is a serious issue. Exposure to ultraviolet radiation and photo-oxidation in combination with wind, wave action and abrasion, degrade plastic fragments into micro and nanosized particles. These particles pass with relative ease through wastewater filters, making their recovery impossible when they reach the sea. Here, corrosion, high temperatures, waves, wind, ultraviolet radiation, and microbial action, continue the slow process of degradation. The small debris remains at the mercy of the currents, floating, going to the bottom, or ending up on the beaches. In fact, most of the seabed all over the world and in the Mediterranean sea in particular, is made of plastic, resulting from the waste recovered on the coasts and in the sea. Microplastics (MPs) are defined as particles less than 5 mm in size^2^. MPs do not derive only from larger pieces fragmentation, but are also produced in these dimensions for commercial uses. They can be found in aqueous, terrestrial and aerial environments^3^. Furthermore, there are several reports of microplastics in food^4^, and in particular in seafood, sea salt^5,6^, and in drinking water^7^. Microplastics have also been detected in the gastrointestinal tract of marine animals^8,9^.

Inside tissues, MPs and microparticles are considered as foreign bodies by the host organism and, as such, trigger local immunoreactions. Furthermore, they can act as carriers for other chemicals, such as environmental pollutants or plastic additives, which are known for their harmful effects^10,11^.

Although there are recent reports highlighting public health concerns due to microplastics presence in food, to date there is little data available. A study reports detection of microplastics in the human intestine^12^. There are also reports on microplastics inhalation in humans: this seems to be an important route of diffusion. However, to date microplastics have never been reported within human placentas.

In this study, we investigated, for the first time, the presence of microparticles and microplastics in human placentas. Placenta finely regulates foetal to maternal environment and, indirectly, to the external one, acting as a crucial interface via different complex mechanisms^13^. The potential presence of man-made microparticles in this organ may harm the delicate response of differentiation between self and non-self^14^, with a series of related consequences that need to be defined.

In this light, we performed a Raman Microspectroscopy analysis on digested samples of placenta collected from six consenting patients with uneventful pregnancies, to investigate the presence of microplastics and microparticles.

## METHODS

### Experimental design

This was a pilot observational descriptive preclinical study, with prospective and unicentric open cohort. It was approved by the Ethical Committee Lazio 1 (Protocol N. 352/CE Lazio 1; March 31^th^, 2020), and it was carried out in full accordance with ethical principles, including The Code of Ethics of the World Medical Association (Declaration of Helsinki) for experiments involving humans^15^. To participate to this study, six selected consenting patients signed an informed consent, which included donation of placentas. The experimental design of the study is sketched in Figure 1.

**Figure 1.**
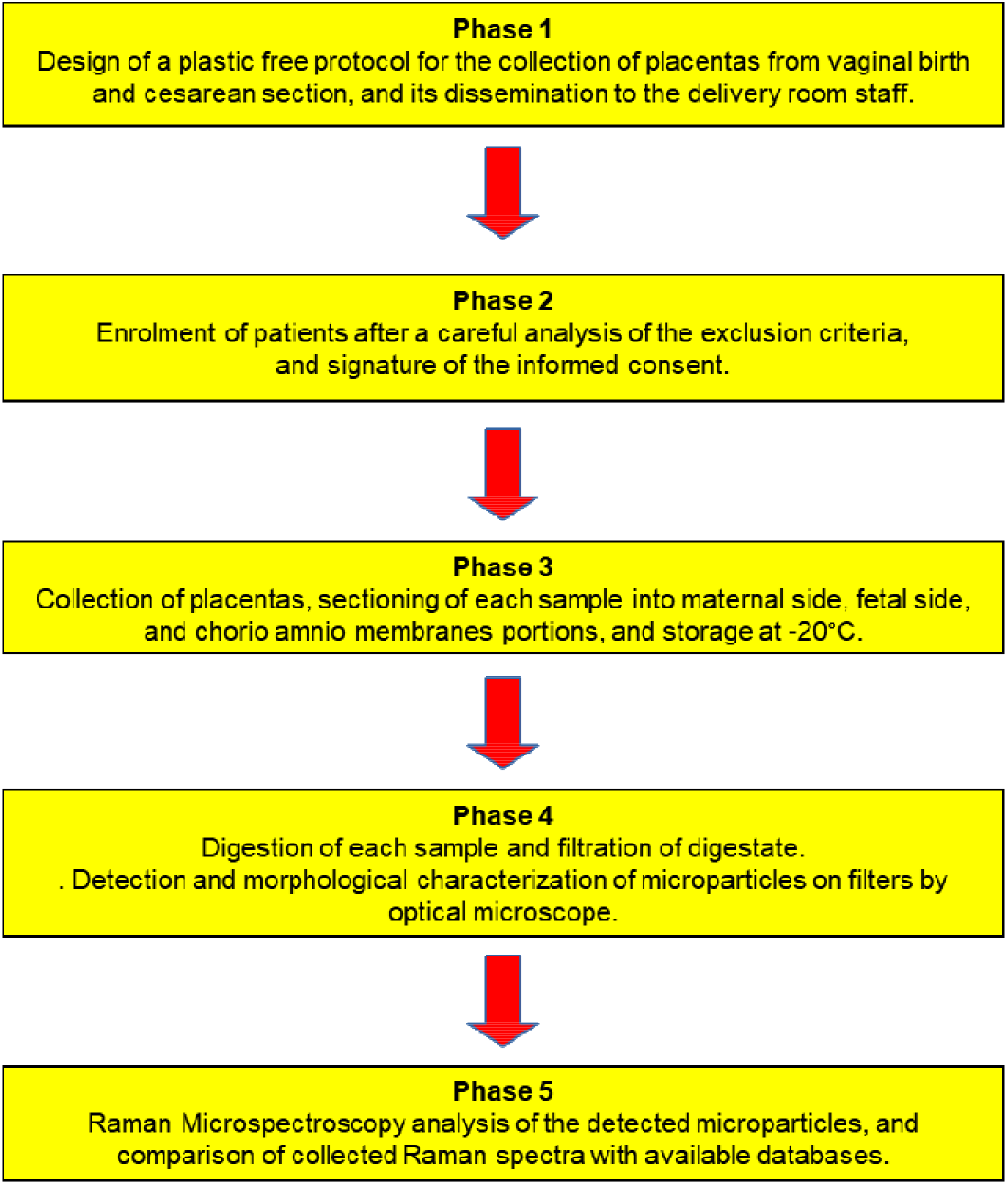
Design of the study.

### Enrolment of patients and placentas collection

All recruited women were healthy, at term of pregnancy. Exclusion criteria were:

- peculiar diets prescribed for any particular medical condition, 4 weeks before delivery;
- diarrhoea or constipation, two weeks before delivery;
- antibiotics intake, two weeks before delivery;
- assumption of drugs affecting intestinal reabsorption (such as activated charcoal, or cholestyramine), two weeks before delivery;
- diagnosis of gastrointestinal disease (such as ulcerative colitis, or Crohn’s disease), cancer, organ transplantation, HIV, or other severe pathologies that needed medical treatment;
- invasive or abrasive dental treatment, two weeks before delivery;
- participation to a clinical study, four weeks before delivery
- alcohol abuse (defined as a >10 score in the Alcohol Use Disorders Identification Test).

In order to get information on the chemicals taken by patients the week before delivery, women were asked to fill a questionnaire to record their food consumption (omnivorous, vegetarian, vegan, with no diet restriction), with particular attention to seafood, food sealed in plastic containers/films, beverages in plastic bottles, carbonated drinks, alcoholic drinks, chewing gums containing microplastics. Moreover, patients were asked to take note of the use of toothpastes and cosmetics containing microplastics or synthetic polymers, and cigarette smoking.

All six women had a vaginal delivery, at the Department of Obstetrics and Gynaecology of San Giovanni Calibita Fatebenefratelli Hospital, Isola Tiberina, Roma (Italy). All placentas were collected according to a protocol specially designed to be plastic-free, with a special focus on avoiding contaminations from plastic fibres or particles. Obstetricians and midwives used cotton gloves to assist women in labour. In the delivery room, only cotton towels were used to cover patients’ beds; graduate bags to estimate postpartum blood loss were not used during delivery, but they were brought in the delivery room only after birth, when umbilical cord was already clamped and cut with metal clippers, avoiding contact with plastic material. After birth, placentas were deposed onto a metal container and immediately taken to the Laboratory of Pathological Anatomy, San Giovanni Calibita Fatebenefratelli Hospital, Isola T\iberina, Roma (Italy). Pathologist, wearing cotton gloves and using metal scalpels, collected from each placenta, portions (mean weight: 23.3 ± 5.7 g) taken from maternal side, foetal side, and chorioamniotic membranes. All samples, strictly anonymous, were labelled with number codes and stored in glass bottles with metal lids at −20°C with no further treatment. By expedited refrigerated transport, samples were shipped to the Laboratory of Vibrational Spectroscopy, Department of Life and Environmental Sciences, Università Politecnica delle Marche (Ancona, Italy).

### Extraction of microparticles from placenta samples

The extraction of microparticles from the portions of placenta, collected at San Giovanni Calibita Fatebenefratelli Hospital (Rome, Italy) and their analysis by Raman Microspectroscopy were performed at the Laboratory of Vibrational Spectroscopy, Department of Life and Environmental Sciences, Università Politecnica delle Marche (Ancona, Italy). To prevent plastic contamination, cotton laboratory coats, face masks and single-use latex gloves were worn during sample handling, preparation of samples and during the entire experiment. Work surfaces were thoroughly washed with 70% ethanol prior starting all procedures. All liquids (deionised water for cleaning and for preparation of KOH solution) were filtered through 1.6 μm-pore-size filter membrane (Whatman GF/A). Glassware and instruments, including scissors, tweezers and scalpels, were washed using dishwashing liquid, rinsed with deionised water and finally rinsed with 1.6 μm-filtered deionised water. Since the experiments were conducted without the use of the laminar flow hood, the plastic fibres found in the samples were not considered in the results.

Microparticles’ isolation from placenta samples was performed modifying the protocols from two previous works^16,17^. Samples were weighed and placed in a glass container cleaned as previously explained. A 10% KOH solution was prepared using 1.6 μm-filtered deionised water and KOH tablets (Sigma-Aldrich). This solution was added to each jar in a ratio with the sample of 1:8 (w/v). The containers were then sealed and incubated at room temperature for 7 days.

Digestates were then filtered through 1.6 μm-pore-size filter membrane (Whatman GF/A) using a vacuum pump connected to a filter funnel. The filter papers were dried at room temperature and stored in glass Petri dishes until visual identification and spectroscopic characterization of particles. Three procedural blanks, obtained following the same procedure above described, but without placenta samples and maintained close to the samples during their manipulation, were tested to monitor and correct potential contaminations^16^.

### Analysis of microparticles by Raman Microspectroscopy

The analysis of microparticles found in the placenta samples was performed by a Raman XploRA Nano Microspectrometer (Horiba Scientific). The following protocol was adopted: (1) to highlight the presence of microparticles (<20 μm), filter membranes were inspected by visible light using a ×10 objective (Olympus MPLAN10x/0.25); (2) the detected microparticles were first morphologically characterized by a ×100 objective (Olympus MPLAN100x/0.90), and (3) then directly analyzed on the filter by Raman Microspectroscopy (spectral range 160-2000 cm^−1^, 785 nm laser diode, 600 lines per mm grating). The spectra were dispersed onto a 16-bit dynamic range Peltier cooled CCD detector; the spectrometer was calibrated to the 520.7 cm^−1^ line of silicon prior to spectral acquisition.

Raw Raman spectra were submitted to polynomial baseline correction and vector normalization, in order to reduce noise and enhance spectrum quality (Labspec 6 software, Horiba Scientific). The collected Raman spectra were compared with those reported in the SLOPP Library of Microplastics (Spectral Library of Plastic Particles^18^) and in the spectral library of the KnowItAll software (Bio-Rad Laboratories, Inc.). Similarities of more than 80 of Hit Quality Index (HQI) were considered satisfactory.

All data generated or analysed during this study are included in this published article.

## RESULTS

From each placenta, three portions (one from maternal side, one from foetal side and one from chorioamniotic membranes, for a total of eighteen pieces) were collected and processed for the subsequent analysis by Raman Microspectroscopy, to verify the presence of microplastics and, more in general, of microparticles similar to man-made products. As described in the Methods section, strict precautions were taken to prevent contaminations; no microparticles were detected on the filters of the blank procedural samples. In total, 12 microparticles, characterized as microplastics and other man-made materials, were detected in the placentas of 4 out of the 6 enrolled patients. In particular, 5 microparticles were found in the foetal side portions, 4 in the maternal side portions, and 3 in the chorioamniotic membranes. All the analyzed microparticles were pigmented; pigments are usually added to polymers in order to colour plastic products, and are added also to coloured paints and coatings, which are ubiquitous as microplastics^19^.

A retrospective analysis based on Raman spectral information and data reported in literature was performed to define the nature of these microparticles. Firstly, the collected Raman spectra were compared with those stored in the spectral library of the KnowItAll software (Bio-Rad Laboratories, Inc.). Due to the presence of pigments, in many cases, collected Raman spectra resulted mainly due to the signals of the pigment itself^19,20^. It is known that Raman scattering is more sensitive to the chemical functional groups of pigments, which cover with their signals the entire Raman spectrum, than to the polymeric matrix^21^. In these cases, the KnowItAll software allows to identify the pigments contained in the microparticles. By matching the results from the KnowItAll software with the information obtained by consulting the European Chemical Agency (ECHA^22^), it was possible to accurately identify the commercial name, chemical formula, IUPAC name and Color Index Constitution Number of all pigments. Then, in order to uncover the identity of the polymer matrix of the detected microparticles, the collected Raman spectra were also compared with those reported in the SLOPP library of microplastics (Spectral Library of Plastic Particles^18^). Only for three microparticles, it was possible to unveil the signals of the polymer matrix in the spectrum. Table 1 reports the morphological and chemical features of the detected microparticles and relative pigments, together with information regarding the placenta portion in which they were found.

**Table 1.**
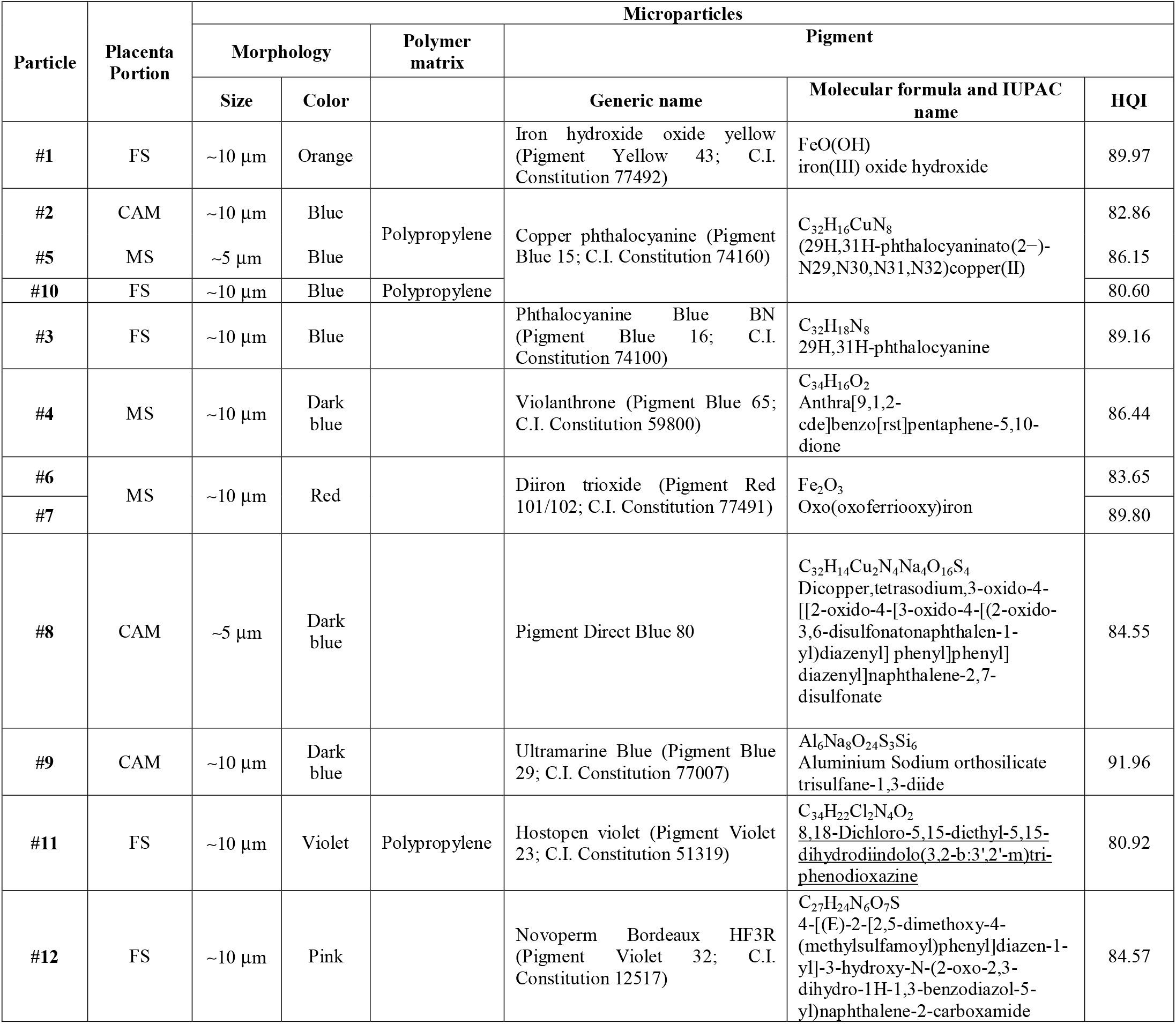
Morphological and chemical features of the detected microparticles and relative pigments, together with information regarding the placenta portion in which they were found (fetal side FS; maternal side MS, and chorio amnio membrane CAM; Hit Quality Index HQI).

The microphotographs of the analyzed microparticles are reported in Figure 2, together with the collected Raman spectra.

**Figure 2.**
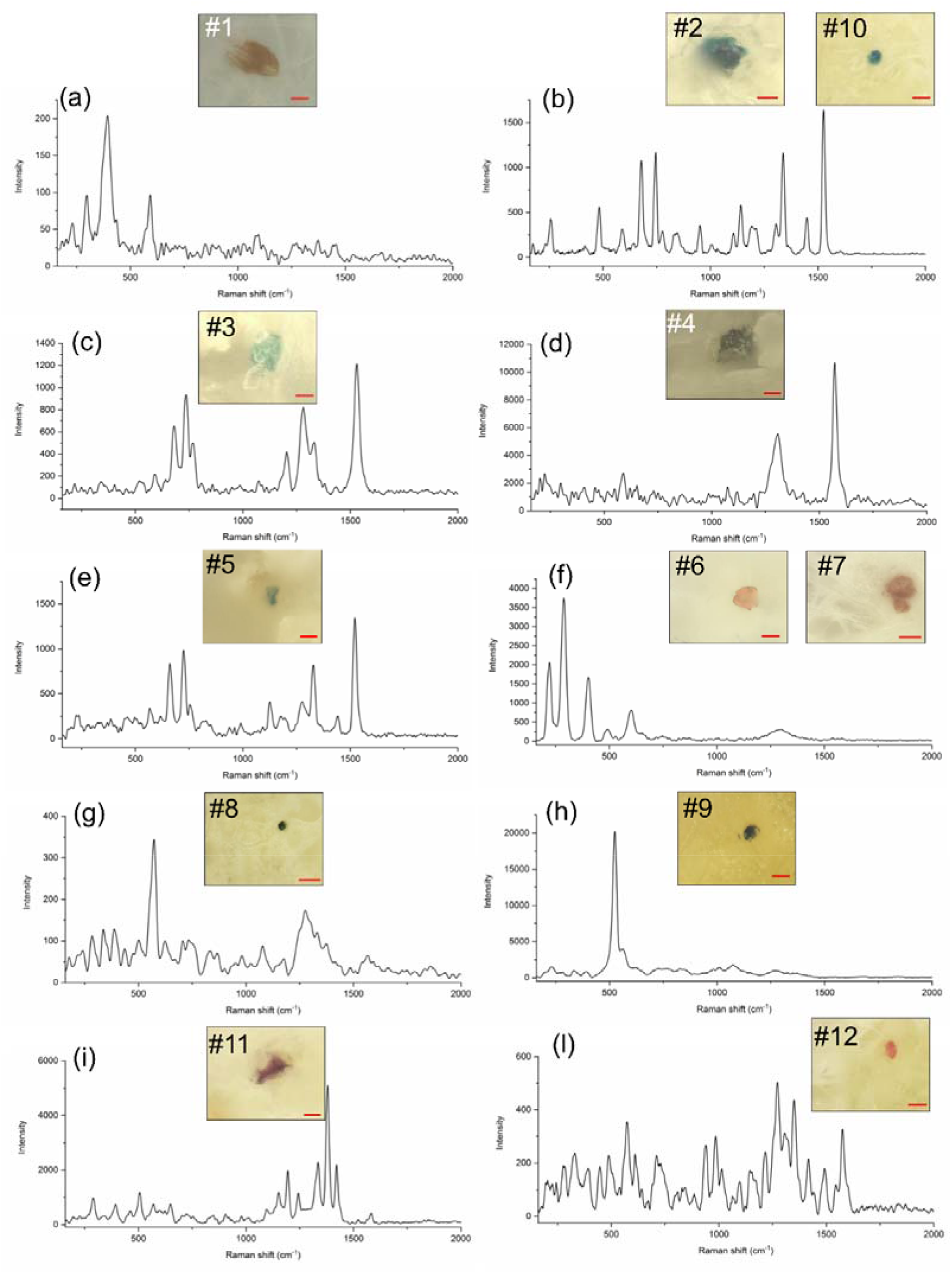
Microphotographs and collected Raman spectra of: (a) Particle #1 (scale bar 5 μm); (b) Particles #2 and #10 (scale bar 5 μm for #2 and 10 μm for #10); (c) Particle #3 (scale bar 5 μm); (d) Particle #4 (scale bar 5 μm); (e) Particle #5 (scale bar 5 μm); (f) Particles #6 and #7 (scale bar 10 μm for #6 and 5 μm for #7); (g) Particle #8 (scale bar 10 μm); (h) Particle #9 (scale bar 10 μm); (i) Particle #11 (scale bar 5 μm), and (l) Particle #12 (scale bar 10 μm).

The interpretation of the spectral data is discussed below.

**Particle #1** (Figure 2a). The collected Raman spectrum resulted perfectly superimposable to the one of the pigment Iron hydroxide oxide yellow: the two spectra shared the main peak at 396 cm^−1^, related to the vibrations of iron oxides/hydroxides. This pigment is described as powder or particulate, and it is used for coloration of polymers (plastics and rubber) and in a wide variety of cosmetics, such as BB creams and foundations.

**Particles #2 and #10** (Figure 2b). The collected Raman spectra resulted comparable to the one of a polypropylene (PP) blue sample. The Raman spectra of the identified particles shared with the reference spectrum the position of the main peaks, such as the peaks centred at 253 cm^−1^ (wagging of CH_2_ moieties, bending of CH moieties), 397 cm^−1^ (wagging of CH_2_ moieties, bending of CH moieties), 839 cm^−1^ (rocking of CH_2_ and CH_3_ moieties, stretching of CC and C-CH_3_ moieties), 970 cm^−1^ (rocking of CH_3_ moieties), and 1455 cm^−1^ (bending of CH_3_ and CH_2_ moieties), all assigned to PP^23^. The bands at 679 cm^−1^, 1143 cm^−1^, 1340 cm^−1^ and 1527 cm^−1^, common to reference blue polypropylene and sample spectra, are known to be related to Raman signals of blue pigments, mainly based on copper phthalocyanine^24,25^.

**Particle #3** (Figure 2c). The collected Raman spectrum resulted superimposable to the one of the blue pigment phthalocyanine^25^. This chemical is reported to be used in adhesives, coating products, plasters, finger paints, polymers and cosmetics and personal care products.

**Particle #4** (Figure 2d). The collected Raman spectrum resulted superimposable to the one of the pigment violanthrone. This chemical is used especially for textile (cotton/polyester) dyeing, coating products, adhesives, fragrances and air fresheners. The two main peaks composing both reference and sample spectra are those centred at 1573 cm^−1^ (C-C stretching of benzene ring) and 1307 cm^−1^ (in reference spectrum, an additional shoulder at ~1350 cm^−1^ is visible, assigned to C-C stretching and HCC bending).

**Particle #5** (Figure 2e). The collected Raman spectrum resulted perfectly superimposable to the one of the pigment copper phthalocyanine^25^. Hence, differently from the particle #2 and #10, it was not possible unveiling the identity of the polymer matrix. This pigment is reported to be used for staining of plastic materials, made of polyvinylchloride (PVC), low density polyethylene (LDPE), high density polyethylene (HDPE), polypropylene (PP), polyethylene terephthalate (PET). Furthermore, the pigment 74160 is widely used for staining coating products and finger paints.

**Particles #6 and #7** (Figures 2f). The collected Raman spectra resulted perfectly superimposable to the one of the red pigment oxo (oxoferriooxy) iron: the two spectra shared with the reference one the three main peaks at 220, 287 and 401 cm^−1^, typical of iron oxides^26^. The same pigment is reported as Pigment Red 101 and 102, depending on its synthetic or natural origin. This pigment is used as food additive, for coloration of plastics, rubber, textiles and paper.

**Particle #8** (Figure 2g). The collected Raman spectrum resulted superimposable to the one of the pigment Direct Blue 80. This dye is reported to be used for coloration of leather, paper and textiles.

**Particle #9** (Figure 2h). The collected Raman spectrum resulted superimposable to the one of the pigment Ultramarine Blue. This pigment is mainly applied in cosmetics, for example for formulations of soap, lipstick, mascara, eye shadow and other make-up products.

**Particle #11** (Figure 2i). The collected Raman spectrum resulted comparable to the one of a PP purple fibre. The Raman spectrum of the identified particle shared with the reference spectrum all the positions of the main peaks, partly ascribable to PP^23^ (such as the peaks centred at 397 cm^−1^, assigned to the wagging of CH_2_ moieties/bending of CH moieties, and at 1455 cm^−1^, assigned to the bending of CH_3_ and CH_2_ moieties), but mainly ascribable to the violet pigment (such as the bands centred at 1193 cm^−1^, 1335 cm^−1^ and 1381 cm^−1^)^25^.

**Particle #12** (Figure 2l). The collected Raman spectrum resulted superimposable to the one of the pink pigment Novoperm Bordeaux HF3R^25^. The Raman spectrum of this monoazopigment shared with the sample spectrum the main peaks centred at 731 cm^−1^, 961 cm^−1^, 1219 cm^−1^, 1280 cm^−1^, 1360 cm^−1^, and 1580 cm^−1^. This pigment is reported to be used to permanently coat and protect wood surfaces, in photographic chemicals, inks and toners, given its high solvent resistance and good heat stability.

## DISCUSSION

In this study, we reported, for the first time, the presence of man-made microparticles and MPs in human placentas. The analysis of portions of maternal side, foetal side and chorioamniotic membranes of human placentas revealed the presence of 12 pigmented microparticles, compatible with microplastics and other man-made materials, in the placentas of 4 women out of the total of the 6 analyzed. In particular, 5 microparticles were found in the foetal side, 4 in the maternal side and 3 in the chorioamniotic membranes, indicating that these microparticles, once internalized, can colonize placenta tissues at all levels (Figure 3).

**Figure 3.**
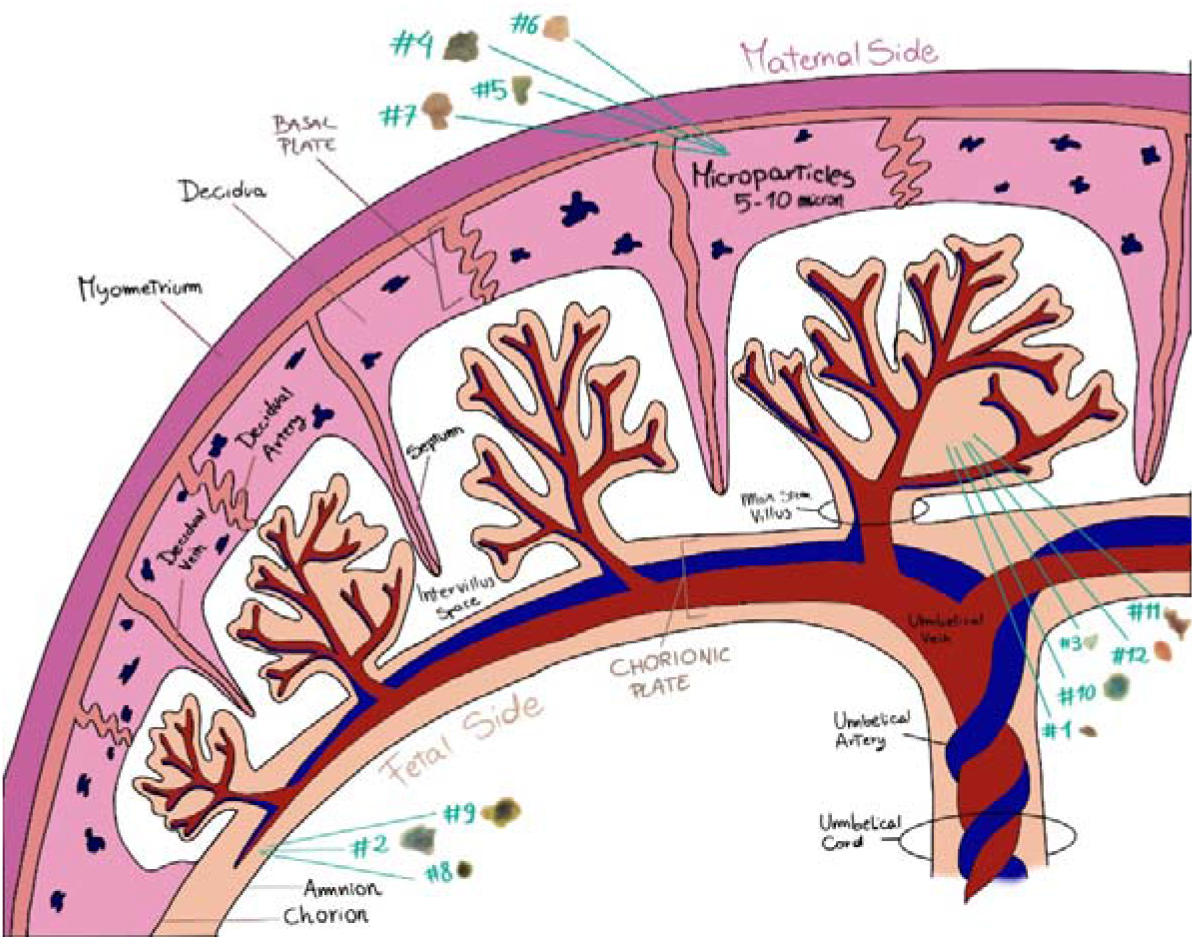
The figure illustrates the twelve microparticles that we found in the analyzed placentas and described in figure 1. They are located in the placental portion in which they were found.

The identified microparticles were differentiated between stained microplastics (particles #2, #10 and #11, all attributable to polypropylene) and paint/coating/dye microparticles, in which the polymer matrix had lower amount (particles #1, #3-9, and #12) ^19^. All the microparticles were ~10 μm in size, except for two that were smaller (~5 μm): these dimensions are compatible with a possible transportation by bloodstream. Previous analyses of 5-10 μm particles, by Electron Microscopy coupled whit X-ray microprobe, revealed the presence of microparticles as foreign bodies in human internal organs^27^.

Microparticles and MPs may access the bloodstream and reach placenta from the gastrointestinal tract (GIT)^28^, from the maternal respiratory system (Figure 4A-B-C-D), or both, by M cells-mediated endocytosis mechanisms, or paracellular transport. It is known that the fraction of inhaled particles, with less than 2.5 μm, is largely retained in the lungs, but can pass through respiratory barriers^29^. The microparticles isolated in the present study have dimensions of 5-10 μm, making it plausible that they were removed from the respiratory cilia, once internalized by inhalation; in effect, the most probable transport routes for nanoparticles is the diffusion through cellular membranes, while particles with dimensions of 10-20 μm may reach internal organs mostly by mechanisms of particle uptake and translocation, as described for the internalization from the GIT^30^. GIT persorption is described as the translocation of particles into the circulatory system of the GIT through gaps in the epithelium of the villus tips; it is expected to represent the major uptake route for microparticles. Uptake and subsequent translocation to secondary target organs depend on several factors, including hydrophobicity, surface charge, surface functionalization and the associated protein corona, and particle size. The uptake and translocation to secondary target organs of microparticles were associated with inflammatory responses in the surrounding tissues, such as the immune activation of macrophages and the production of cytokines^31^.

**Figure 4 A-B-C-D.**
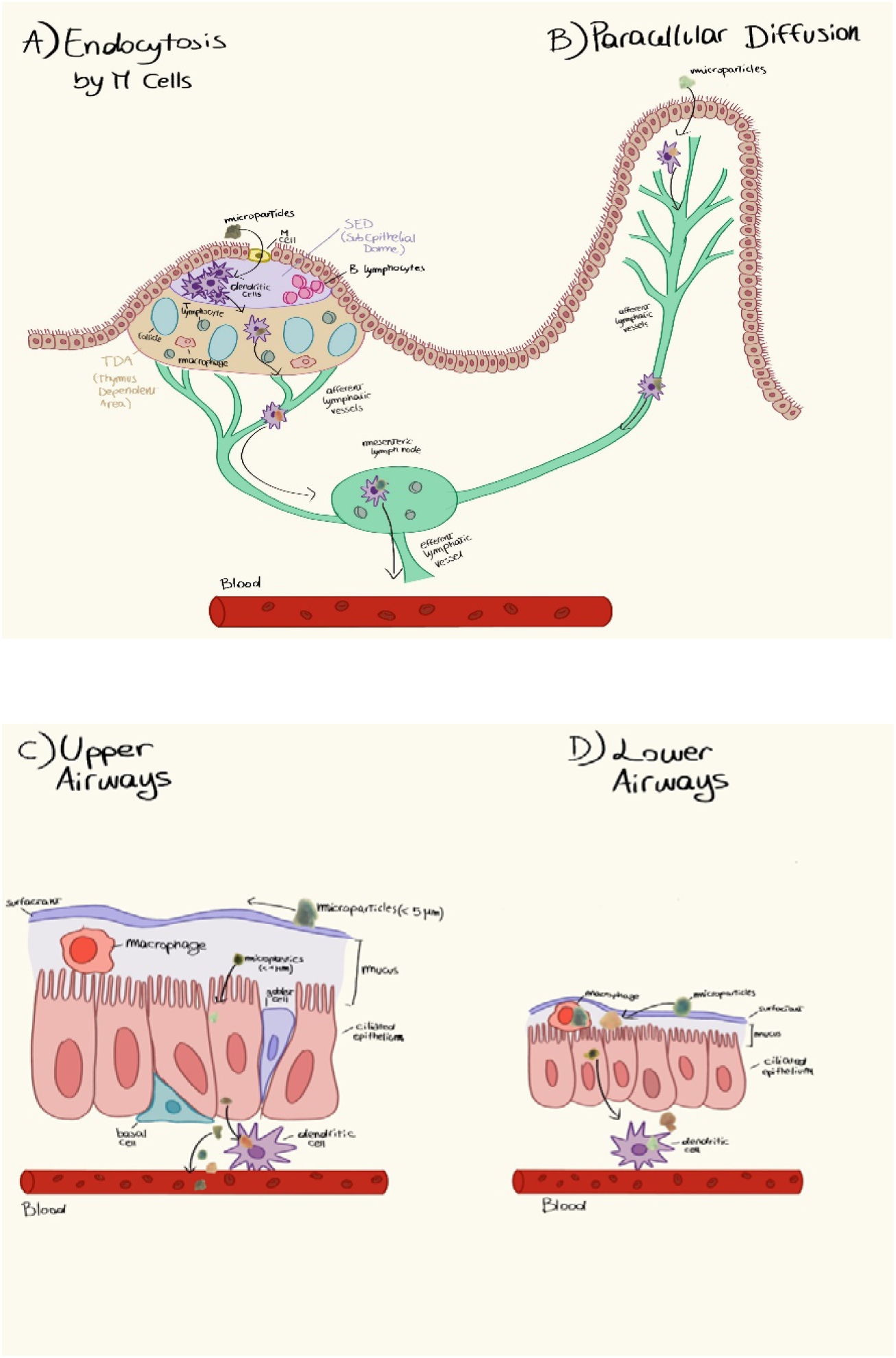
Hypothetical mechanisms by which microplastics penetrate human tissues. (A) Endocytosis by M cells. At the level of the Peyer's Patched, below the mucous gut, MPs ingested with food can be uptaken by endocytosis from the M cells, transported across the epithelium into the subepithelial dome where they encounter dendritic cells, which in turn transport them through the lymphatic circulation, from where they reach the blood. (B) Paracellular Diffusion. MPs could penetrate through the intestinal lumen from loose junctions. This phenomenon could partially explain why some inflammatory states, which increase loose junctions favour intestinal passage. Once the intestinal lumen has been crossed, the MPs are collected by the dendritic cells and transported in the lymphatic and subsequently in the systemic circulation. (C) Upper airways, At the level of the upper respiratory tract the mucus is thicker and allows a successful clearance of the foreign bodies particles, in addition, the mechanical movement of ciliated epithelium and the presence of surfactant prevents smaller particles from spreading through the epithelium and reach the circulation. (D) Lower airways, In the lower respiratory tract the mucus layer is thinner, thus facilitating the diffusion of particles which, thanks to their particular aerodynamic shape, are able to reach this part of the respiratory tract. Once penetrated, the MPs can spread into the general circulation by cellular uptake or diffusion. (*Modified from: Mowat, A. Anatomical basis of tolerance and immunity to intestinal antigens. Nat Rev Immunol **3,** 331–341 (2003). https://doi.org/10.1038/nri1057. And Ruge, C. A.; Kirch, J.; Lehr, C. M. Pulmonary drug delivery: From generating aerosols to overcoming biological barriers-therapeutic possibilities and technological challenges. Lancet. Respir. Med. 2013, 1(5), 402−413.)*

Once microparticles have reached the maternal surface of the placenta (Figure 3), they can invade the tissue in depth by several transport mechanisms, both active and passive, that are not clearly understood yet^32^. The transplacental passage of 5-10 μm size microparticles may depend on the different physiological conditions and genetic characteristics of placenta: this may explain, together with the diverse food habits and lifestyle of patients, the absence of microparticles in 2 of the 6 analyzed placentas and the different localization and characteristics of the particles identified in the present study. It is known that a great variability exists in the expression and function of placental drug transporters, both within human populations (interindividual variability) and also during gestation (intraindividual variability)^33^. We suppose that this variability exists also in relation to the mechanism of particles’ internalization.

The presence of microparticles in the placenta tissue requires to reconsider the immunological mechanism of self-tolerance. Placenta represents the interface between the foetus and the environment^13^. Embryos and foetuses must continuously adapt to the maternal environment and, indirectly, to the external one, by a series of complex responses. An important part of this series of responses consists in differentiate self and non-self^14^, a mechanism that may be perturbed by the presence of microparticles and MPs. It is in fact reported that, once internalized, MPs may accumulate and exert localized toxicity by inducing and/or enhancing immune responses and, hence, potentially reducing the defence mechanisms against pathogens and altering the utilization of energy stores^10^.

Potentially, in placenta, MPs, and in general microparticles, may alter several cellular regulating pathways, such as immunity mechanisms during pregnancy, growth-factor signalling during and after implantation, functions of atypical chemokine receptors governing maternal-foetal communication, signalling between the embryo and the uterus, and trafficking of uterine dendritic cells, natural killer cells, T cells and macrophages during normal pregnancy. All these effects may lead to adverse pregnancy outcomes^34^. Three of the particles identified in the present study (particles #2, #10, and #12) resulted polypropylene (PP). It is known that polymers used in plastic products have cytotoxic effects. For example, the toxicity of PP particles appears related to their size: smaller PP particles may provide more surface area to disturb cell growth. Moreover, it was observed that, when administered as a powder, PP particles, neither smaller nor larger, were cytotoxic, while PP particles dispersed in medium have potentially greater toxicity. The administration of PP particles of dimensions of 5-10 μm resulted in inducing murine macrophage cells to increase IL-6 secretion, suggesting that small PP particles may mimic potential pathogens^35^.

A crucial problem related to microplastics is their potential release of chemicals, which can cause severe damages to cells. In fact, plastic debris has shown to contain various contaminants, including micromolecular substances such as chemicals and monomers. Some of these substances, such as bisphenol A, phthalates and some of the brominated flame retardants, are endocrine disruptors, known to adversely affect human health upon exposure via ingestion and inhalation^36^. It is reported that low concentrations of bisphenol A can affect cell proliferation in human placental first trimester trophoblasts, downregulating mRNA expression of VEGF and causing an abnormal placental development^37^. Moreover, phthalates have been found in human urine and blood samples; they are considered responsible of several effects in animals and humans, such as impairment of pubertal development, male and female reproductive health, pregnancy outcomes and respiratory health^38^.

In conclusion, this is the first study revealing the presence of man-made microparticles in human placenta, shedding new light on the level of human exposure to microplastics and microparticles in general. The dimensions of the detected particles are consistent with the known mechanisms of particle uptake and translocation, described for other internalization routes and yet to be clarified in this organ. Due to the crucial role of placenta in hosting the foetus and in acting as an interface between the latter and the external environment, the presence of exogenous and potentially harmful particles is matter of great concern, for the possible consequences on pregnancy outcomes. Further studies need to be performed to increase the number of enrolled patients. Moreover, we are planning to investigate if microparticles are in the intracellular or extracellular compartment of tissues, moving from a digestion-based protocol to a histology-based one. Finally, further analyses will be necessary to assess if the presence of these particles in human placenta may trigger immune responses or determine the release of toxic contaminants, resulting harmful for pregnancy.

## Acknowledgements

We would like to thank all San Giovanni Calibita Fatebenefratelli Hospital staff for their collaboration during the study and placentas collection.

## Author contributions

A.R. and E.G. designed the study; C.S., P.C., V.N., O.C., F.P., M.C.A.R., F.B., S.D., E.D.A. and D.R. performed experiments; A.R., A.S., C.S., P.C., V.N., O.C., F.P., M.C.A.R., F.B., S.D., E.D.A., D.R. and E.G. analysed and interpreted data; A.R., E.G. M.M. and A.S. drafted the manuscript;

## Correspondence and requests for materials

should be addressed to A.S.

